# Efficiency of genome-wide association study in open-pollinated populations

**DOI:** 10.1101/050955

**Authors:** José Marcelo Soriano Viana, Gabriel Borges Mundim, Fabyano Fonseca e Silva, Antonio Augusto Franco Garcia

**Author notes:** Corresponding author: José Marcelo Soriano Viana. Federal University of Viçosa, Department of General Biology, 36570-900, Viçosa, MG, Brazil. Federal University of Viçosa, Department of General Biology, 36570-900, Viçosa, MG, Brazil. Federal University of Viçosa, Department of Animal Science, 36570-900, Viçosa, MG, Brazil. ESALQ - University of São Paulo, Department of Genetics, 13418-900, Piracicaba, SP, Brazil.

## Abstract

Genome-wide association studies (GWAS) with plant species have employed inbred lines panels. Thus, to our knowledge, no information is available on theory and efficiency of GWAS in open-pollinated populations. Our objectives are to present quantitative genetics theory for GWAS, evaluate the relative efficiency of GWAS in non-inbred and inbred populations and in an inbred lines panel, and assess factors affecting GWAS, such as linkage disequilibrium (LD), sample size, and quantitative trait locus (QTL) heritability. Fifty samples of 400 individuals from populations with LD were simulated. Individuals were genotyped for 10,000 single nucleotide polymorphisms (SNPs) and phenotyped for traits with different degrees of dominance controlled by 10 QTLs and 90 minor genes. The average SNP density was 0.1 centiMorgan and the trait heritabilities were 0.4 and 0.8. We assessed GWAS efficiency based on the power of QTL detection, number of false-positive associations, bias in the estimated QTL position, and range of the significant SNPs for the same QTL. When the LD between a QTL and one or more SNPs is restricted to markers very close to or within the QTL, GWAS in open-pollinated populations can be highly efficient, depending mainly on QTL heritability and sample size. GWAS achieved the highest power of QTL detection, the smallest number of false-positive associations, and the lowest bias in the estimated QTL position for the inbred lines panel correcting for population structure. Under low QTL heritability and reduced sample size, GWAS is ineffective for non-inbred and inbred populations and for inbred lines panel.

## INTRODUCTION

Association mapping is a high-resolution method for mapping quantitative trait locus (QTL) based on linkage disequilibrium (LD) (Yu and Buckler 2006). Linkage disequilibrium is commonly defined as the non-random association of alleles at two loci carried on the same gamete, caused by their shared history of mutation and recombination (Weir 2008; Flint-Garcia *et al.* 2003). Association mapping has been successful in detecting genes controlling human diseases and quantitative traits in plant and animal species (Pearson and Manolio 2008; Zhu *et al.* 2008; Barendse *et al.* 2007). There are two main association mapping strategies: the candidate gene approach, which focuses on polymorphisms in specific genes that control the traits of interest, and the genome-wide association study (GWAS), which surveys the entire genome for polymorphisms associated with complex traits (Rafalski 2010).

With the advent of high-throughput genotyping and sequencing technologies, breeders have used GWAS to identify genes underlying quantitative trait variation. Compared to QTL mapping, which has statistical precision in the range of 10 to 30 centiMorgans (cM) (confidence interval or highest probability density interval), the main advantage of GWAS is a more precise identification of candidate genes (Zhu *et al.* 2008). Another advantage is the use of a breeding population instead of one derived by crossing two inbred or pure lines (Flint-Garcia *et al.* 2005). However, as highlighted by Weir (2010), the efficiency of GWAS is considerably affected by relatedness and population structure, which can generate spurious association between unlinked marker and QTL. Rafalski (2010) emphasized that the choices of population (due to the degree of LD and genotypic variation), marker density, and sample size are crucial decisions for achieving greater power of QTL detection. Ingvarsson and Street (2011) discussed the influence of population size, extent of LD, trait heritability (precision of phenotyping), and population structure on GWAS efficiency, highlighting that studies with plant species should greatly increase population size to detect QTLs with lower effect (heritability of 1-2%).

Yu *et al.* (2006) proposed a mixed model approach for GWAS analysis called the Q + K (or QK) method, where Q and K are the population structure and kinship matrices, respectively. This method has provided the best results and greatly has improved the control of both type I and type II error rates compared with other methods. Stich and Melchinger (2009) and Yang *et al.* (2010) compared GWAS methods based on simulated and field data. Based on type I error control and power of QTL detection, they concluded that the mixed model approach using only the kinship matrix (K model) to correct for relatedness was more efficient than the approaches controlling only population structure (Q model) and both population structure and relatedness because the spurious associations could not be completely controlled by population structure. Based on simulated inbred lines panel, Bernardo (2013) demonstrated that his models G and QG were superior to the K and QK, respectively, where G indicates a model that uses genome-wide markers to account for QTLs on background chromosomes. The new approach showed a better balance between power of QTL detection and false discovery rate (FDR).

Recently, many instances of GWAS have been published with plant species, including barley, sorghum, wheat, rice, sugarcane, soybean and particularly maize (Ingvarsson and Street 2011). Pace *et al.* (2015) carried out a GWAS with 384 maize inbred lines evaluated for 22 seedling root architecture traits and genotyped with 681,257 single nucleotide polymorphisms (SNPs). They identified 268 marker-trait associations. Some of these SNPs were located within or near (less than one kilo base pairs) to candidate genes involved in root development at the seedling stage. Thirunavukkarasu *et al.* (2014) evaluated 240 elite inbred lines of subtropical maize under water stress and used a set of 29,619 high-quality SNPs. The GWAS identified 50 SNPs consistently associated with agronomic traits related to functional traits that could lead to drought tolerance. Thirty-one of the SNPs detected were situated near drought-tolerance genes. Schaefer and Bernardo (2013) used GWAS on a collection of 284 historical maize inbred lines and 39,166 SNPs and identified 19 QTLs for flowering time, 13 for kernel composition, and 22 for disease resistance. However, only two candidate genes were suggested: one regulating days to anthesis and one regulating oil concentration. Additionally, several QTL hot spots (chromosome regions with previously mapped QTLs) were also identified, affecting days to anthesis (four), days to silking (two), and resistance to northern corn leaf blight (four) and Goss’s wilt and blight (one).

Genome-wide association studies with plant species have employed inbred lines panels. Thus, to our knowledge, no information is available on theory and efficiency of GWAS in open-pollinated populations. Our objectives are to present quantitative genetics theory for GWAS, to evaluate the relative efficiency of GWAS in non-inbred and inbred populations and in an inbred lines panel, and to assess factors that affect GWAS, such as LD, sample size, and QTL heritability.

## MATERIALS AND METHODS

### Quantitative genetics theory for GWAS in open-pollinated populations

Consider a biallelic QTL (alleles **B**/**b**) and a SNP (alleles **C**/**c**) located in the same chromosome, and a population (generation 0) of an open-pollinated species. Assuming LD, the joint gamete and joint genotype probabilities in the population are presented by Viana et al. (2016). The QTL genotypic values are G_**BB**_ = m_b_ + a_b_, G_**Bb**_ = m_b_ + d_b_, and G_**bb**_ = m_b_ - a_b_, where m_b_ is the mean of the genotypic values of the homozygotes, a_b_ is the deviation between the genotypic value of the homozygote of higher expression and m_b_, and d_b_ is the dominance deviation (the deviation between the genotypic value of the heterozygote and m_b_). The average genotypic values of individuals with the genotypes **CC**, **Cc**, and **cc** are

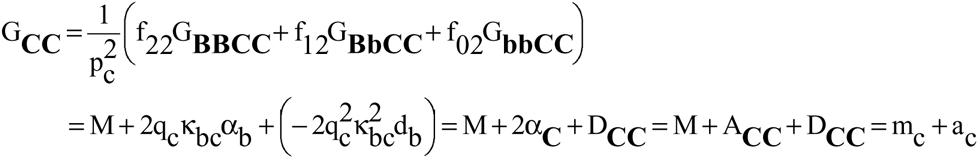

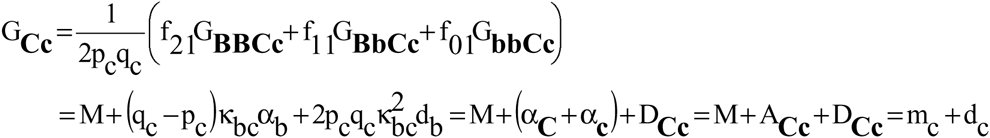

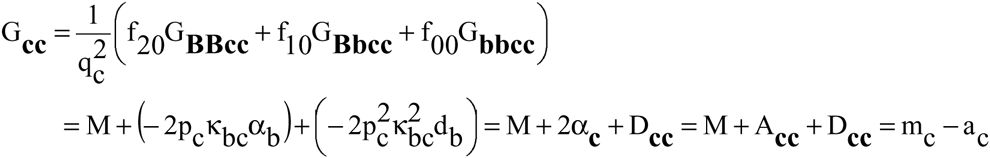

where p is the frequency of the major allele (**B** or **C**), q = 1 - p is the frequency of the minor allele (**b** or **c**), f_ij_ is the probability of the individual with i and j copies of the allele **B** of the QTL and allele **C** of the SNP (i, j = 2, 1, or 0) (for simplicity, we omitted the superscript (0) - for generation 0 - in all parameters that depend on the LD measure of generation −1), M = m_b_ + (p_b_ - q_b_)a_b_ + 2p_b_ q_b_ d_b_ is the population mean, 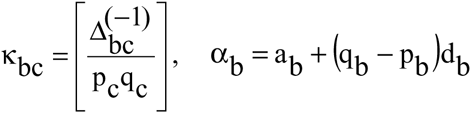 αb = a_b_ + (q_b_ - p_b_)d_b_ is the average effect of a gene substitution, α_**C**_ = q_c_ k_bc_ α_b_ and α_**c**_ = −p_c_ k_bc_ α_b_ are the average effects of the SNP alleles, and A and D are the SNP additive and dominance values. 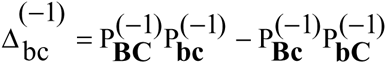 is the measure of LD in the gametic pool of generation −1 (Kempthorne 1957), where P^(−1)^ indicates a joint gamete probability. Another common measure of LD is the square of the correlation between the values of the alleles at the two loci 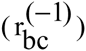 in the gametic pool of generation −1 (Hill and Robertson 1968). Note that 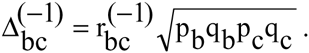 The average effect of substituting the allele **C** for **c** is α_SNP_ = α_**C**_ - α_**c**_ = k_bc_ α_b_. The dominance deviation for the SNP is 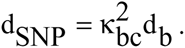 The other SNP parameters are m_c_ = M + (q_c_-p_c_)α_SNP_-(1-2p_c_q_c_)d_SNP_, a_c_ = α_SNP_ -(q_c_ −p_c_)d_SNP_, and d_c_ = d_SNP_.

Assuming no QTL in LD with the SNP, G_**CC**_ = G_**Cc**_ = G_**cc**_ = M. Thus, the identification of the QTL can be based on testing the hypothesis that there is no difference between these genotypic means (based on analysis of variance). Assuming thousands of SNPs, it is necessary to employ a Bonferroni-type procedure to control the type I error when there are multiple-comparisons, as that proposed by Benjamini and Hochberg (1995). Note that α_SNP_ = a_c_ + (q_c_ - p_c_)d_SNP_, where 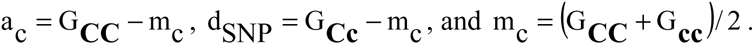

Alternatively, the QTL identification can be done by testing that there is no relationship between the genotypic values for the individuals **CC**, **Cc**, and **cc** with the number of copies of one SNP allele. The parameters of the additive-dominance model can be derived by fitting the model 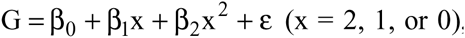, where G is the QTL genotypic value. The model can be expressed as y_(9 x 1)_ = X_(9 x 3)_.β_(3 x 1)_ + error vector_(9 x1)_, where y is the vector of QTL genotypic values, conditional on the SNP genotype, X is the incidence matrix, and β is the parameter vector. The matrix of genotype probabilities is P_(9 _x_ 9)_ = diagonal{f_ij_}. Thus, for the complete model or a reduced model, 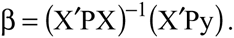. The parameters for the complete model are

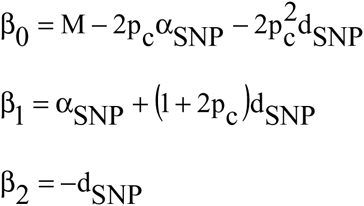

The alternative regression model is 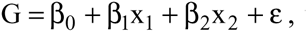, where x1 = 1, 0, or −1 if the individual is **CC**, **Cc**, or **cc**, and x_2_ = 0 or 1 if the individual is homozygous or heterozygous, respectively. Fitting the complete model, β_0_ = m_c_, β_1_ = a_c_, and β_2_ = d_SNP_. Assuming no QTL in LD with the SNP, β_1_ = β_2_ = 0 and β_0_ = M, regardless of the model. Fitting the additive model, 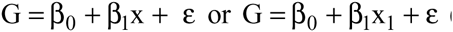 (no dominance), β_1_ = β_SNP_.

If there are two QTLs (alleles **B**/**b** and **E**/**e**) in LD with the SNP (alleles **C**/c), it can be demonstrated that (Viana *et al.* 2016)

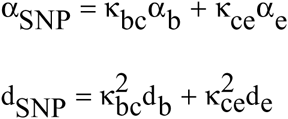

where 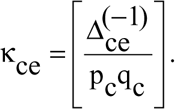Thus, the average effect of a SNP substitution (and the SNP additive value) is proportional to the measure of LD and to the average effect of a gene substitution for each QTL that is in LD with the marker, and the SNP dominance deviation (and the SNP dominance value) is proportional to the square of the LD value and to the dominance deviation for each QTL that is in LD with the marker.

If there is population structure, this must be corrected in the GWAS to avoid spurious associations due to admixture LD. For simplicity, consider two subpopulations in Hardy-Weinberg equilibrium and a SNP (alleles **C**/**c**) and a QTL (alleles **B**/**b**) unlinked, in linkage equilibrium in both subpopulations. Assuming that p and q are the allelic frequencies in one subpopulation and r and s are the allelic frequencies in the other subpopulation, the average genotypic value of individuals **CC**, **Cc**, and **cc** are

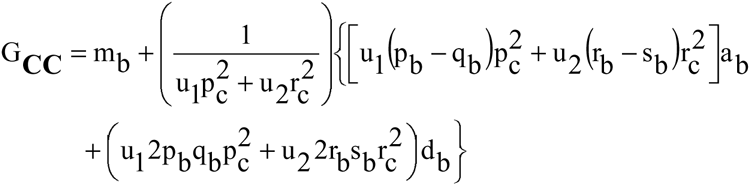

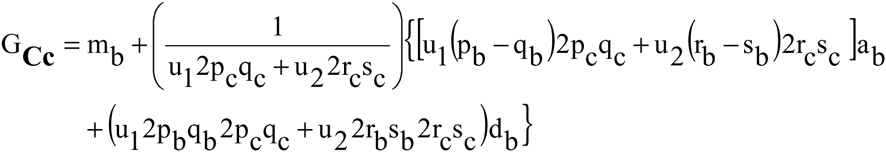

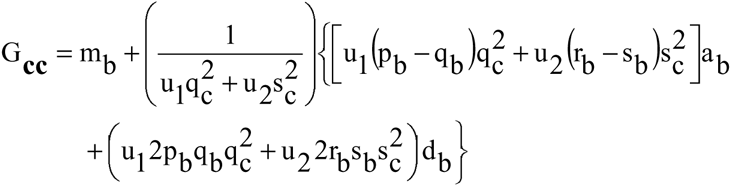

where u_1_ and u_2_ are the proportions of individuals from subpopulations 1 and 2 (probabilities of an individual belongs to subpopulations 1 and 2). Only if there is no population structure (u_1_ = 1 or 0), G_**CC**_ = G_**Cc**_ = G_**cc**_ = M (and β_1_ = β_2_ = 0 and β_0_ = M).

### Quantitative genetics theory for GWAS with inbred lines panel

In general, the inbred lines in a panel represent the genetic variability for the traits being assessed. Therefore, an inbred lines panel includes inbreds from distinct populations or heterotic groups. Consider again a QTL (alleles **B**/**b**) and a SNP (alleles **C**/**c**) located in the same chromosome, and that they are in LD in a population (generation 0). Assuming n (n→∞) generations of selfing, the (limits of the) probabilities of the inbreds (recombinant inbred lines; RILs) are (for simplicity, we omitted again the superscript (0)-for generation 0 - in all parameters that depend on the LD measure of generation −1)

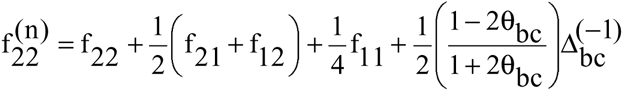

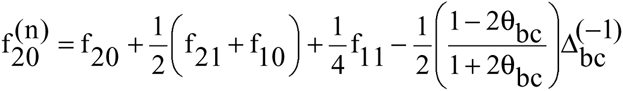

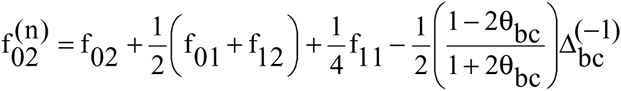

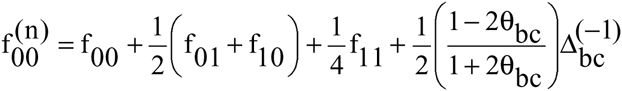

where θ_b_ is the frequency of recombinant gametes. The haplotypes are 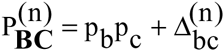, 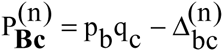, 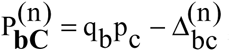, and 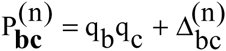, where 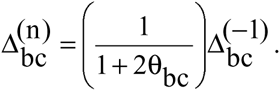 Thus, if there is crossing over (0 < θ_bc_ ≤ 0.5), the LD in this inbred population is lower than the LD in generation −1. If the SNP and QTL are completely linked (θ_bc_ = 0), the LD in the inbred population is the same LD in generation −1. The maximum decrease is 50%, achieved with θ_bc_ = 0.5. Compared with the LD in generation 0, the LD in generation n is 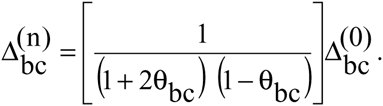 Thus, the maximum decrease is 12%, achieved with θ_bc_ = 0.25. In contrast, after n generations of random crosses 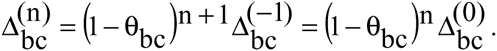 Thus, if 0 < θ_bc_ ≤ 0.5, the maximum decrease is 100% since 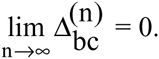 For the inbreds sampled from a population, we have

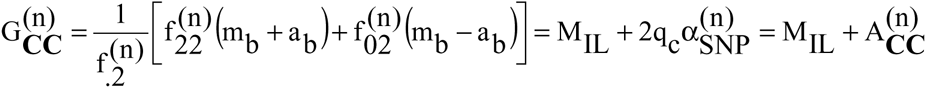

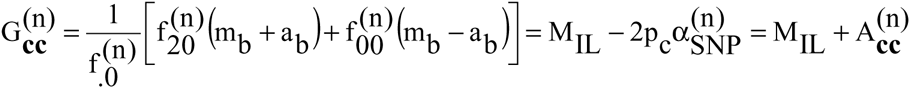

where M_IL_ =m_b_ +(p_b_ −q_b_)a_b_ is the inbred population mean, 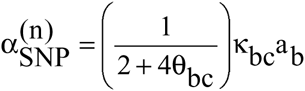 SNP average effect of allele substitution in the inbred population, and A is the SNP additive value for an inbred line. Assuming no QTL in LD with the SNP, 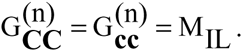 Note that 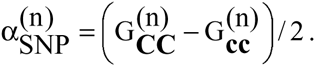

The haplotypes of an inbred lines panel including inbreds from N populations are 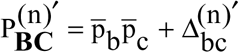, 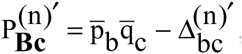, 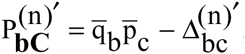, and 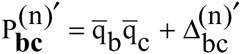, where 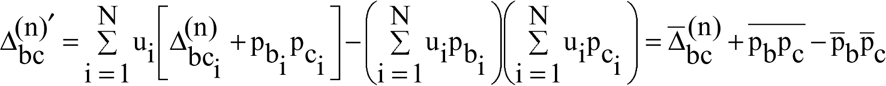 and u_i_ is the probability of an inbred line belonging to population i. Because this function is too complex to interpret, the analysis of the LD value in an inbred lines panel relative to the LD in the inbreds from each population is presented using the simulated data.

Due to population structure, associations involving unlinked SNP and QTL in linkage equilibrium in the non-inbred populations can be declared. For simplicity, assume an inbred lines panel with inbreds from two populations where an unlinked pair of SNP (alleles **C**/**c**) and QTL (alleles **B**/**b**) is in linkage equilibrium. Let and u_1_ and u_2_ be the proportions of inbreds from these populations. Assuming that p and q are the allelic frequencies in one population, that r and s are the allelic frequencies in the other population, and that p ≠ q or r ≠ s,

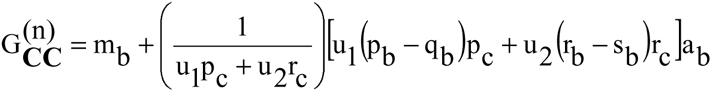

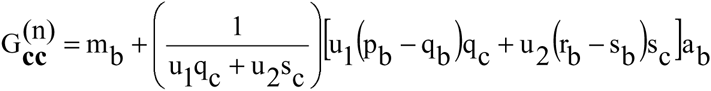

If there is no population structure (u_1_ = 1 or 0), 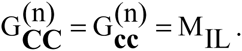

## Simulation

We simulated 50 samples of populations with LD using the software *REALbreeding* (Viana *et al.* 2016, 2013; Azevedo *et al.* 2015). This software has been developed by the first author using the program *REALbasic 2009*. Population 1, generation 10r, is the advanced generation of a composite of two populations in linkage equilibrium (population 1, generation 0) obtained after 10 generations of random crosses, assuming a sample size of 400 individuals. Population 1, generations 10s and 10r10s, were obtained from Population 1, generation 0, assuming 10 generations of selfing and 10 generations of random crosses followed by 10 generations of selfing, respectively, assuming sample sizes of 100 and 400, respectively. Populations 2, 3, and 4, generation 10s, are also inbred populations (10 generations of selfing) derived from composites of two populations, also assuming a sample size of 100. The parents of populations 2 and 3 were assumed to be non-improved and improved populations, respectively. An improved population was defined as having frequencies of favorable genes greater than 0.5, while a non-improved population was defined as having frequencies less than 0.5. A composite is a Hardy-Weinberg equilibrium population with LD for only linked markers and genes. In the case of a composite of two populations in linkage equilibrium, 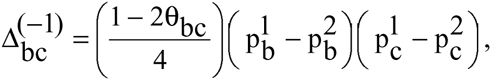, where the indices 1 and 2 refer to the parental populations.

Based on our input, *REALbreeding* randomly distributed 10,000 SNPs, 10 QTLs (of higher effect) and 90 minor genes (QTLs of lower effect) in 10 chromosomes (1,000 SNPs and 10 genes by chromosome). The average SNP density was 0.1 cM. The genes were distributed in the regions covered by the SNPs. Four, three, two, and one QTLs were inserted in chromosomes 1, 5, 9, and 10, respectively. We also specified one SNP within each QTL and a minimum distance between linked QTLs of 10 cM. To allow *REALbreeding* to compute the phenotypic value for each genotyped individual, we informed the minimum and maximum genotypic values for homozygotes, proportion between the parameter *a*for a QTL and the parameter *a* for a minor gene (a_QTL_/a_mg_), degree of dominance ((d/a)i, i = 1,…, 100), direction of dominance, and broad sense heritability. *REALbreeding* saves two main files, one with the marker genotypes and another with the additive, dominance, and phenotypic values (non-inbred populations) or the genotypic and phenotypic values (inbred populations). The true additive and dominance genetic values or genotypic values are computed from the population gene frequencies (random values), LD values, average effects of gene substitution or *a* deviations, and dominance deviations. The phenotypic values are computed from the true population mean, additive and dominance values or genotypic values, and from error effects sampled from a normal distribution. The error variance is computed from the broad sense heritability.

We simulated three popcorn traits. The minimum and maximum genotypic values of homozygotes for grain yield, expansion volume, and days to maturity were 30 and 180 g per plant, 15 and 65 mL/g, and 100 and 170 days, respectively. We defined positive dominance for grain yield (0 < (d/a)_i_ ≤ 1.2), bidirectional dominance for expansion volume (−1.2 ≤ (d/a)_i_ ≤ 1.2), and no dominance for days to maturity ((d/a)_i_ = 0). The broad sense heritabilities were 0.4 and 0.8. These values can be associated with individual and progeny assessment, respectively. Assuming a_QTL_/a_mg_ = 10, each QTL explained approximately 4 and 8% of the phenotypic variance for heritabilities of 0.4 and 0.8. The GWAS was performed in population 1, generations 10r and 10r10s, and in the inbred lines panel obtained from inbreds of the populations 1 through 4, generation 0 (generations 10s). To assess the influence of the sample size on the GWAS efficiency, we considered sample sizes of 400 and 200. Thus, we used 100 or 50 inbreds from populations 1 through 4 to generate the inbred lines panel. To assess the influence of the QTL heritability on the GWAS efficiency, we converted four QTLs (QTLs 3, 7, 8, and 10 on chromosomes 1, 5, 9, and 10, respectively) to minor genes and assumed QTL heritability of 12% (for trait heritability of 0.7). Then, the GWAS was performed on population 1, generation 10r.

### Statistical analyses

The analyses of LD and association were performed with the software *PowerMarker* (Liu and Muse 2005) for open-pollinated populations, and *Tassel* (Bradbury *et al.* 2007) for the inbred lines panel and RILs. Because there is no relationship between the inbred lines, the GWAS with the inbred lines panel was based on the general linear model, correcting for population structure (Q model). For the population structure analysis, we used *Structure* software (Falush *et al.* 2003) and fitted the admixture model with correlated allelic frequencies and the no admixture model with independent allelic frequencies. The number of SNPs, sample size, burn-in period, and number of MCMC (Monte Carlo Markov chain) replications were 100 (10 random SNPs by chromosome), 400 (simulation 1), 10,000, and 40,000, respectively. The number of populations assumed (K) ranged from 1 to 7, and the most probable K value was determined based on the inferred plateau method (Viana *et al.* 2013). We used Benjamini-Hochberg FDR of 5 and 1% to control the type I error (Benjamini and Hochberg 1995).

To classify each significant association as true or false, we used a program developed in *REALbasic 2009* by the first author. The classification criterion was based on the difference between the position of the SNP and the position of a true QTL (candidate gene). If the difference was less than or equal to 2.5 cM (Yu *et al.* 2008), the association was classified as true. The GWAS efficiency was assessed based on the power of QTL detection (probability of rejecting H_0_ when H_0_ is false; control of type II error), number of false-positive associations (control of type I error), bias in the estimated QTL position (precision of mapping), and range of the significant SNPs for the same QTL (Li *et al.* 2010).

### Data availability

*REALbreeding* is available upon request. The data set is available at https://dx.doi.org/10.6084/m9.figshare.3201838.v1. Supplemental file S1 contains detailed description of all data files (SNP and QTL positions, SNP genotypes, and phenotypic values). Data citation: Viana, José Marcelo; Mundim, Gabriel Borges; Fonseca e Silva, Fabyano; Augusto Franco Garcia, Antonio (2016): Efficiency of genome-wide association study in open-pollinated populations. figshare. https://dx.doi.org/10.6084/m9.figshare.3201838.v1

## RESULTS

The results for assessing the efficiency of GWAS in open-pollinated populations refer to population 1, generation 10r. In generation 0, the degree of LD is so high that several significant associations are observed along the length of a chromosome with one or more QTLs or in one or more large chromosome regions (Figure 1). These several significant associations are not false-positive (at least most of them). This is due to the degree of LD and presence of QTL. Even assuming a FDR of 1%, it is worthless for the identification of candidate genes to infer that there are one or more QTLs in a chromosome region spanning 20 cM. When the LD between a QTL and one or more markers is restricted to SNPs very close to or within the QTL, the analysis can be highly efficient, depending mainly on the QTL effect and sample size. Assuming a QTL heritability of 8% and sample size 400 (simulation 1), the significant associations for expansion volume observed in chromosome 1 evidenced five QTLs with a FDR of 5% or four QTLs with a FDR of 1% (Figure 1). This implies in a QTL detection power of 100%. Three of the four true QTLs (candidate genes) were identified by SNPs located within the QTL while one was identified by five or four SNPs in a region spanning approximately 2.0 or 1.7 cM, respectively, depending on the FDR. The significant associations at a FDR of 5 or 1% for SNPs 223 (at position 21.7 cM), 243 (at position 23.3), 245 (at position 23.4 cM), and 252 (at position 23.7 cM) are attributable to their LD with QTL 2. The absolute LD values are 0.1488, 0.1494, 0.1747, and 0.1416, respectively (with highly significant P values according to the chi-square test). The significant association at a FDR of 5% for SNP 627 (at position 61.8 cM) is not a false-positive, since it is in LD with QTL 2 (|Δ| = 0.0366, chi-square test P value = 3.22E-6) and QTL 3 (|Δ| = 0.0302, chi-square test P value = 7.55E-5). Then, the result is interpreted as a fifth QTL.

**Figure 1.**
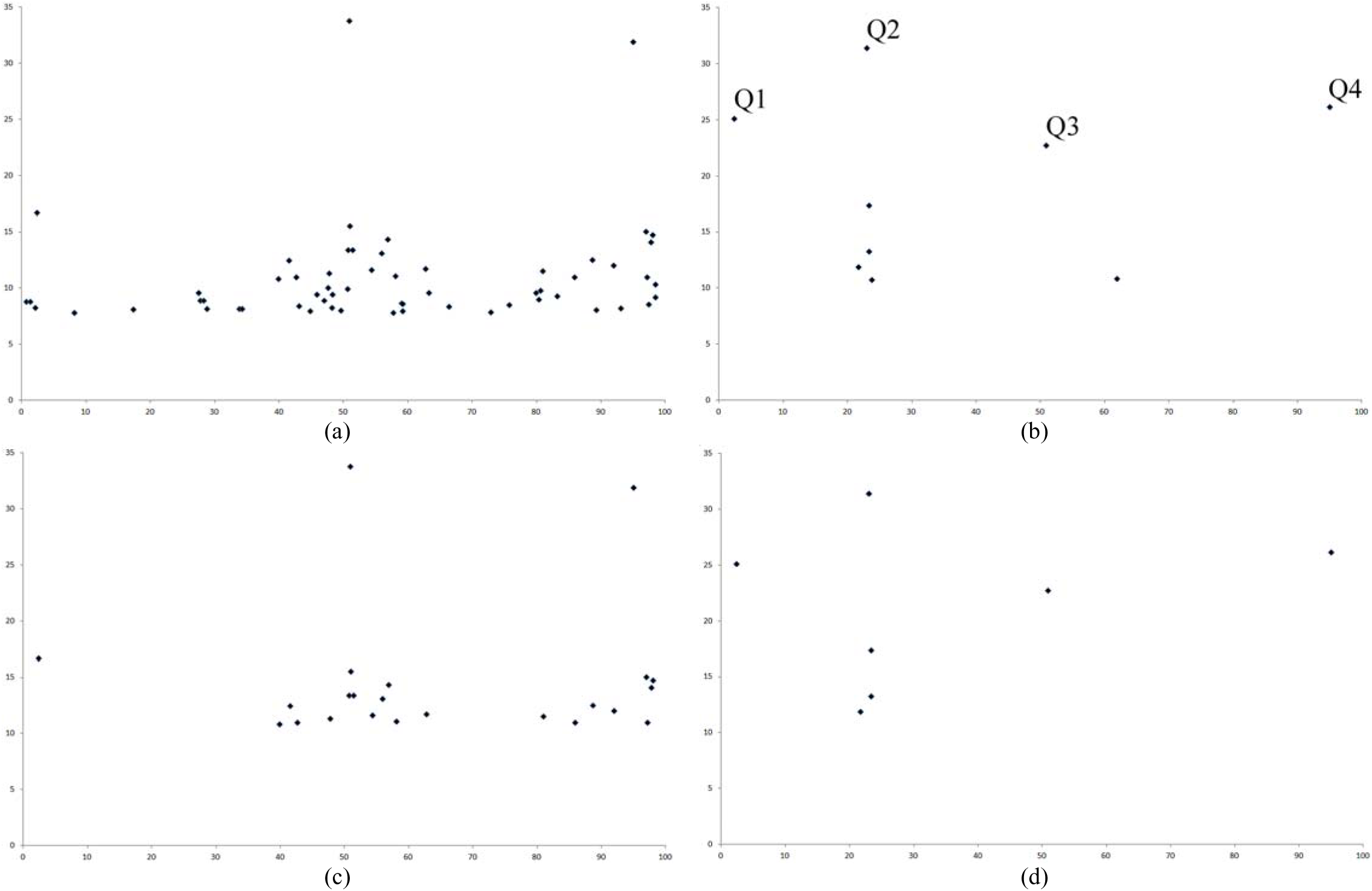
Significant associations at a FDR of 5 (a and b) or 1% (c and d) (F test; Y axe) in chromosome 1 (SNP position (cM); X axe), from the GWAS in population 1, generations 0 (a and c) and 10r (random cross) (b and d), regarding expansion volume, QTL heritability of 8%, and sample size 400 (simulation 1) (Q = QTL).

Only for intermediate to high QTL heritability (8 and 12%) and greater sample size were the results from GWAS clearly different between days to maturity and the other two traits, except for the power of QTL detection (Table 1). The number of significant associations, number of false-positives, bias in QTL position, and average range of chromosome regions with one or more QTLs were greater in the absence of dominance. With a FDR of 5%, the power of QTL detection ranged from 88 to 100% but was associated with a high number of significant associations in chromosomes with one to four QTLs. On average, each true QTL was identified based on two to three (for days to maturity) SNPs, in chromosome regions spanning 0.8 to 2.2 cM. The bias in QTL position ranged from 0.5 to 0.8 cM. Increasing the control of the type I error provided better results and greatly reduced the number of false-positive associations. The power of QTL detection ranged from 75 to 100% and each QTL was identified based on one to two SNPs in chromosome regions spanning 0.4 to 1.1 cM. The bias in QTL position ranged from 0.3 to 0.6 cM.

**Table 1.** Average number of significant associations with a FDR of 5 or 1%, power of QTL detection (%), number of false-positive associations in chromosomes with no QTL and one to four QTLs, bias in the QTL position (cM), and average range for the regions with identified QTL, regarding population 1, generation 10r (random cross), three traits (expansion volume (EV; mL/g), grain yield (GY; g per plant), and days to maturity (DM)), two sample sizes, and three QTL heritabilities^a^

| FDR | Trait | Sample | h<sup>2</sup> | Sig. Assoc. | Power | False+0 | False+1-4 | Bias | Av. range |
| --- | --- | --- | --- | --- | --- | --- | --- | --- | --- |
| 5% | EV | 400 | 4% | 6.7 (0; 22) | 37.2 (0; 80) | 0.6 (0; 5) | 0.9 (0; 8) | 0.21 (0.00; 0.98) | 0.28 (0.00; 2.45) |
|  |  |  | 8% | 32.1 (13; 73) | 88.6 (60; 100) | 3.3 (0; 17) | 7.9 (1; 28) | 0.52 (0.09; 0.83) | 0.84 (0.11; 2.18) |
|  |  |  | 12% | 26.8 (7; 63) | 99.2 (60; 100) | 2.9 (0; 11) | 8.1 (0; 30) | 0.59 (0.00; 0.92) | 1.27 (0.00; 2.74) |
|  |  | 200 | 4% | 0.8 (0; 5) | 5.2 (0; 40) | 0.2 (0; 2) | 0.3 (0; 1) | 0.22 (0.00; 1.76) | 0.14 (0.00; 1.31) |
|  |  |  | 8% | 6.2 (0; 33) | 37.0 (0; 70) | 0.5 (0; 3) | 1.1 (0; 12) | 0.16 (0.00; 0.88) | 0.17 (0.00; 1.42) |
|  |  |  | 12% | 6.0 (2; 20) | 70.8 (20; 100) | 0.5 (0; 4) | 0.8 (0; 5) | 0.18 (0.00; 0.87) | 0.25 (0.00; 1.58) |
|  | GY | 400 | 4% | 5.7 (0; 25) | 34.4 (0; 90) | 0.4 (0; 2) | 0.8 (0; 7) | 0.20 (0.00; 0.85) | 0.18 (0.00; 1.03) |
|  |  |  | 8% | 31.3 (10; 82) | 87.8 (70; 100) | 3.3 (0; 12) | 7.5 (0; 28) | 0.51 (0.00; 0.97) | 0.77 (0.00; 1.54) |
|  |  |  | 12% | 31.6 (8; 74) | 99.6 (80; 100) | 3.6 (0; 16) | 9.8 (0; 43) | 0.68 (0.14; 1.01) | 1.39 (0.12; 2.60) |
|  |  | 200 | 4% | 0.8 (0; 16) | 3.8 (0; 30) | 0.6 (0; 4) | 0.4 (0; 6) | 0.15 (0.00; 1.39) | 0.06 (0.00; 0.94) |
|  |  |  | 8% | 5.9 (0; 18) | 32.6 (10; 80) | 0.8 (0; 8) | 1.1 (0; 7) | 0.16 (0.00; 0.97) | 0.16 (0.00; 1.75) |
|  |  |  | 12% | 7.2 (3; 19) | 74.8 (20; 100) | 0.7 (0; 7) | 1.2 (0; 6) | 0.21 (0.00; 0.75) | 0.30 (0.00; 1.49) |
|  | DM | 400 | 4% | 8.6 (0; 33) | 39.0 (0; 100) | 1.1 (0; 5) | 1.4 (0; 9) | 0.29 (0.00; 0.70) | 0.38 (0.00; 1.64) |
|  |  |  | 8% | 50.7 (12; 119) | 92.8 (70; 100) | 6.5 (0; 21) | 14.9 (3; 48) | 0.65 (0.06; 1.02) | 1.22 (0.07; 2.82) |
|  |  |  | 12% | 50.2 (20; 110) | 100.0 (100; 100) | 6.5 (0; 23) | 18.4 (3; 50) | 0.77 (0.49; 1.05) | 2.02 (0.76; 3.49) |
|  |  | 200 | 4% | 1.0 (0; 7) | 5.0 (0; 30) | 0.5 (0; 2) | 0.5 (0; 3) | 0.17 (0.00; 0.82) | 0.15 (0.00; 2.45) |
|  |  |  | 8% | 6.6 (0; 41) | 34.0 (0; 70) | 0.7 (0; 3) | 1.2 (0; 12) | 0.23 (0.00; 0.91) | 0.27 (0.00; 2.00) |
|  |  |  | 12% | 8.8 (2; 31) | 75.2 (40; 100) | 1.0 (0; 9) | 1.9 (0; 12) | 0.26 (0.00; 0.96) | 0.35 (0.00; 1.38) |
| 1% | EV | 400 | 8% | 15.0 (7; 39) | 76.6 (50; 100) | 0.4 (0; 3) | 2.0 (0; 13) | 0.32 (0.00; 0.75) | 0.41 (0.00; 1.70) |
|  |  |  | 12% | 13.3 (4; 38) | 98.4 (60; 100) | 0.5 (0; 2) | 2.7 (0; 16) | 0.40 (0.00; 0.79) | 0.65 (0.00; 1.92) |
|  | GY | 400 | 8% | 14.8 (6; 39) | 75.4 (60; 100) | 0.3 (0; 3) | 2.1 (0; 10) | 0.33 (0.00; 0.80) | 0.43 (0.00; 1.32) |
|  |  |  | 12% | 15.6 (5; 43) | 98.4 (80; 100) | 0.6 (0; 7) | 3.2 (0; 21) | 0.53 (0.03; 0.81) | 0.82 (0.03; 1.63) |
|  | DM | 400 | 8% | 20.6 (6; 51) | 80.0 (40; 100) | 0.6 (0; 3) | 4.1 (0; 19) | 0.44 (0.00; 0.93) | 0.61 (0.00; 1.99) |
|  |  |  | 12% | 22.3 (8; 49) | 100.0 (100; 100) | 0.9 (0; 6) | 6.5 (0; 21) | 0.58 (0.02; 0.99) | 1.14 (0.03; 2.98) |

<sup>a</sup>the values between parentheses are the minimum and maximum.

Assuming a QTL heritability of 8% and sample size of 200 or a QTL heritability of 4% and sample size of 400, it is better to assume a FDR of 5% to ensure greater power of QTL detection and fewer false-positive associations. However, the power of detection ranged from 33 to 39%, particularly due to the lower QTL effect (Table 1). With lower QTL heritability and reduced sample size, GWAS is ineffective, showing an average power of QTL detection less than or equal to 5%. This scenario does not improve when increasing the FDR to 10% (data not shown). Increasing the QTL heritability to 12% resulted in an increase in the power of QTL detection, particularly when assuming a sample size of 200 individuals (Table 1). There were also increases in bias in the QTL position, range of chromosome regions with an identified QTL, number of false-positives, and number of significant associations in chromosomes with one to two QTLs, mainly with greater sample size. When assuming 200 individuals, the power of QTL detection reached 70-75%, regardless of the trait.

We also provided results for comparing GWAS in open-pollinated population and in an inbred lines panel. An impressive result from GWAS with an inbred lines panel is the efficacy of discarding spurious associations due to population structure (Figure 2). The number of spurious associations in chromosome 3 (no QTL) were reduced from 477 to zero in the analysis of expansion volume assuming a FDR of 1%, QTL heritability of 8%, and sample size 400 (simulation 1). Correcting for population structure decreased the number of significant associations in chromosome 1 (four QTLs) from 464 to 9. This implies a QTL detection power of 100% but with three to five false-positive associations. The population structure analysis evidenced four subpopulations (Figure 3). In general, the efficiency of GWAS was greater with the inbred lines panel (Table 2). The power of QTL detection was higher, and the number of false-positive associations was lower. Furthermore, only SNPs within QTL showed significant associations in general. Also, no differences were observed between the traits and similarly for open-pollinated populations the analysis is ineffective when assuming lower QTL heritability and sample size.

**Figure 2.**
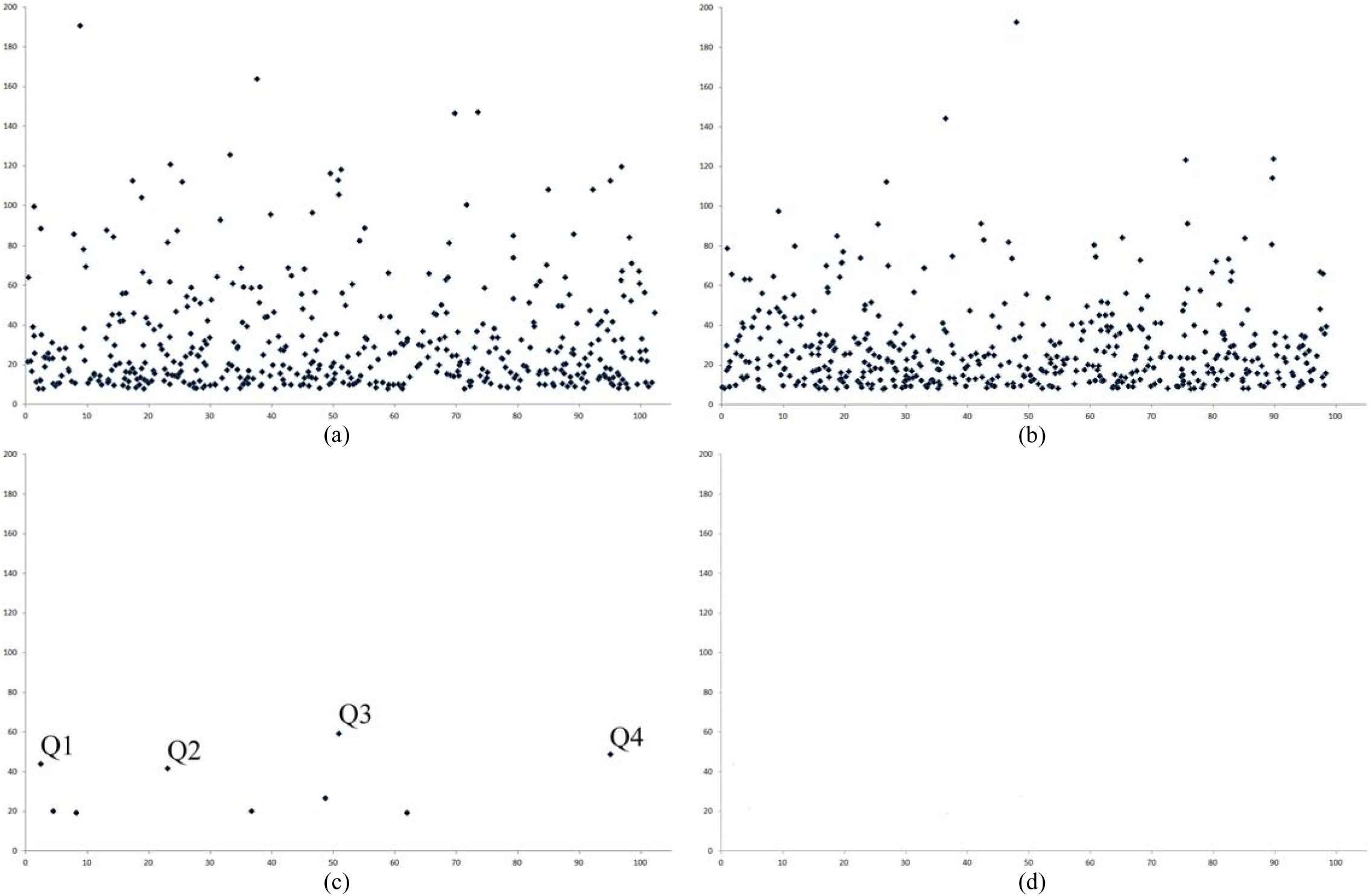
Significant associations at a FDR of 1% (F test; Y axe) in chromosomes 1 and 3 (SNP position (cM); X axe) ignoring (a and b, respectively) and correcting for the population structure (c and d, respectively), from the GWAS in an inbred lines panel regarding expansion volume, QTL heritability of 8%, and sample size 400 (simulation 1) (Q = QTL).

**Figure 3.**
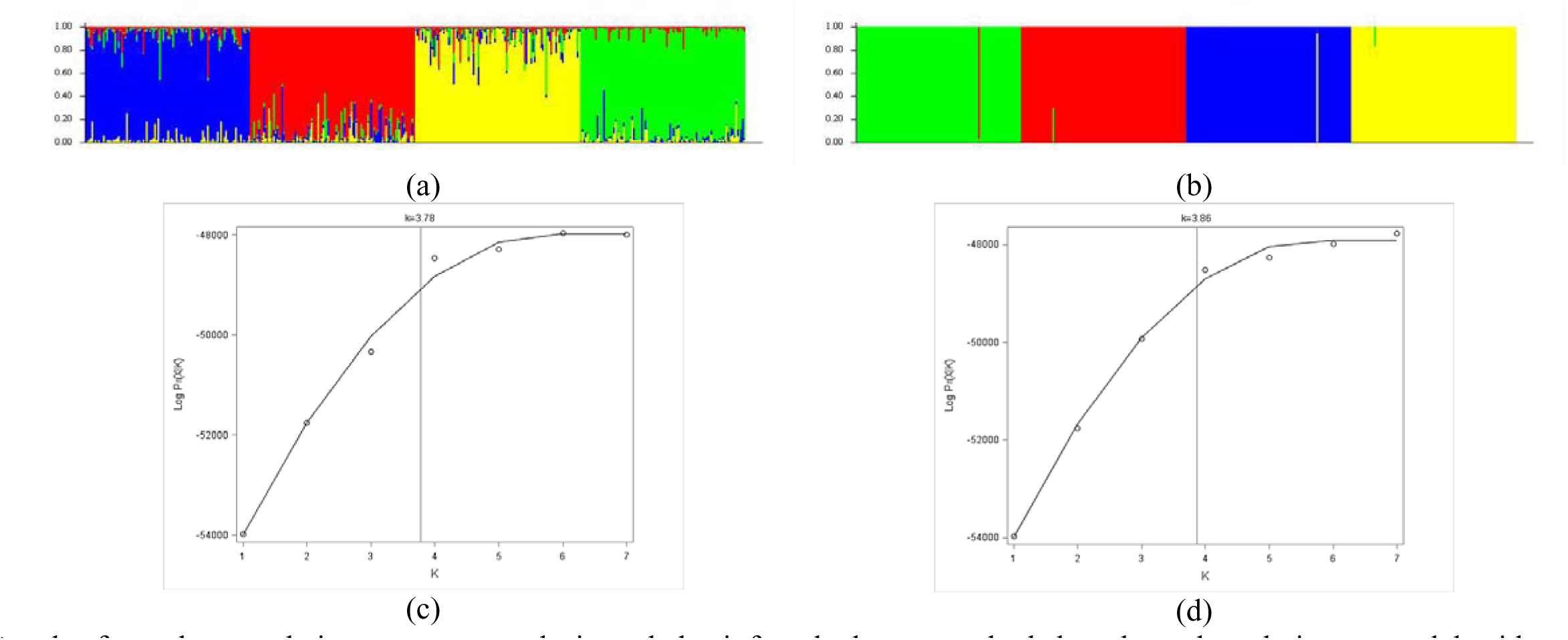
Results from the population structure analysis and the inferred plateau method, based on the admixture model with correlated allelic frequencies (a and c) and the no admixture model with independent allelic frequencies (b and d).

**Table 2.**
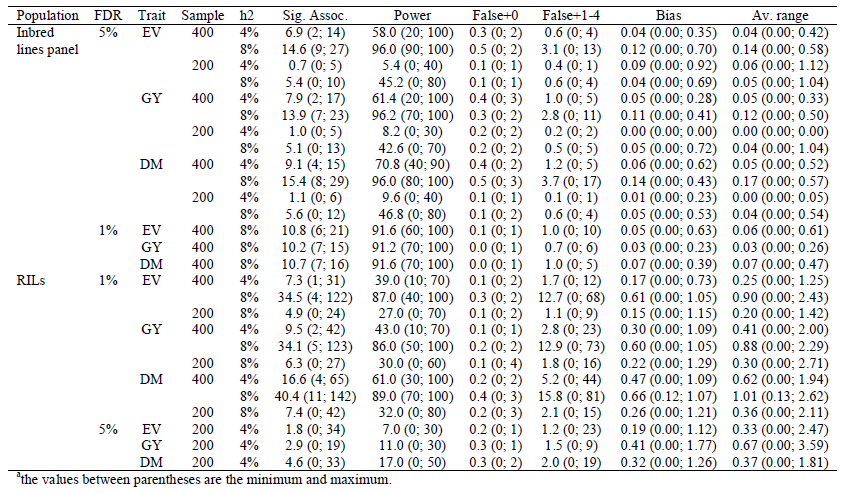
Average number of significant associations with a FDR of 5 or 1%, power of QTL detection (%), number of false-positive associations in chromosomes with no QTL and one to four QTLs, bias in the QTL position (cM), and average range for the regions with identified QTL, regarding an inbred lines panel and RILs from population 1, generation 10r (random cross), three traits (expansion volume (EV; mL/g), grain yield (GY; g per plant), and days to maturity (DM)), two sample sizes, and two QTL heritabilities^a^

The following were indicated by analysis of the parametric LD in the populations and in the inbred lines panel based on a random 10 cM segment of chromosome 1 (100 SNPs): higher LD in population 1, generation 0 (average absolute Δ = 0.0403; 627 values greater than 0.1), lower LD in population 3, generation 0 (average absolute Δ = 0.0203; 48 values greater than 0.1), a slight decrease in the LD with selfing (5-6%), and the lowest LD in the inbred lines panel (average absolute Δ = 0.0249; 8 values greater than 0.1) (Figures 4 and 5). The LD decay due to 10 generations of random crosses was approximately 25%, regardless of the population. For example, the number of absolute LD values greater than 0.1 decreased 60% in population 1, generation 10r.

**Figure 4.**
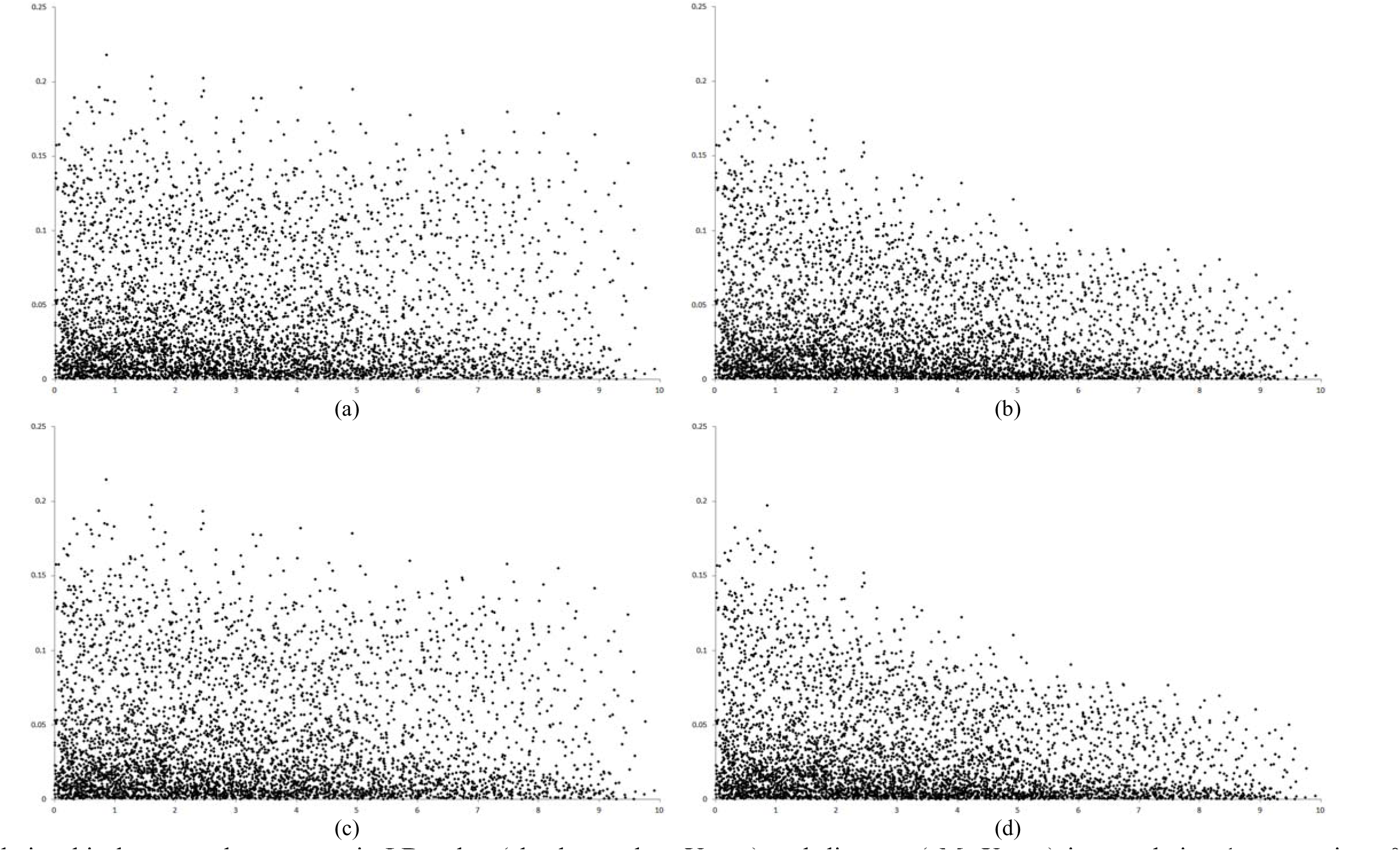
Relationship between the parametric LD value (absolute value; Y axe) and distance (cM; X axe) in population 1, generations 0 (a), 10r (random cross) (b), 10s (selfing) (c), and 10r10s (d), assuming a segment of 10 cM of chromosome 1 (centered on QTL 3).

**Figure 5.**
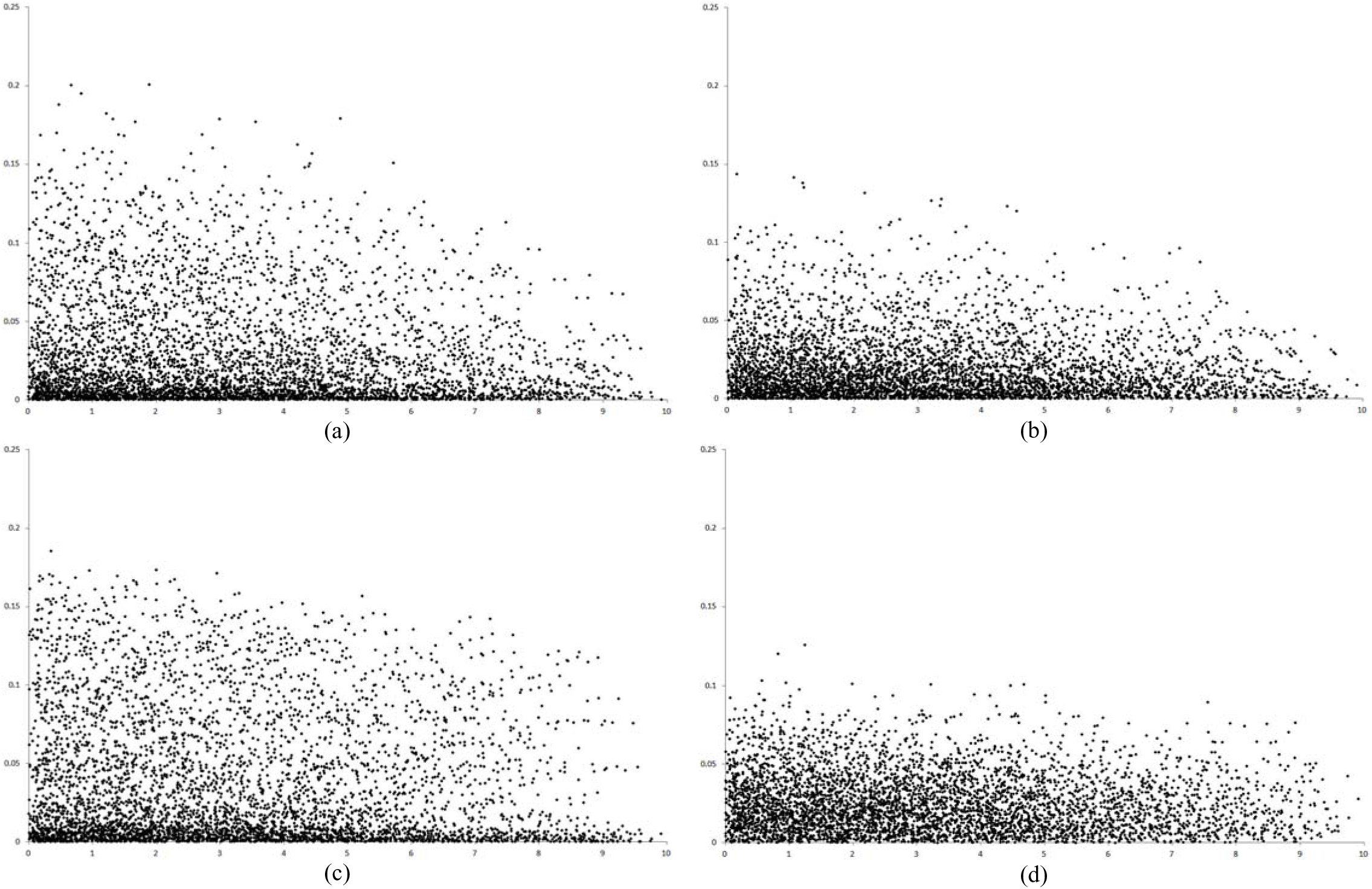
Relationship between the parametric LD value (absolute value; Y axe) and distance (cM; X axe) in populations 2 (a), 3 (b), and 4 (c), generation 10s (selfing), and in the inbred lines panel (d), assuming a segment of 10 cM of chromosome 1 (centered on QTL 3).

Compared to GWAS in population 1, generation 10r, at a FDR of 5%, the GWAS with RILs from population 1, generation 10r (lowest parametric LD among the non-inbred populations), at a FDR of 1%, showed the same power of QTL detection and a high number of significant associations along the length of one or more chromosomes with one to four QTLs (Table 2). As explained, this makes the GWAS ineffective for identifying candidate genes. Compared to GWAS in generation 10r, the lower efficiency of GWAS with RILs for identifying candidate genes (due to a greater number of significant associations in chromosomes with one to four QTLs) can be attributed to higher heritability, due to increase in the genotypic variance for the same error variance, and higher estimated LD. Based on simulation 1, the estimated QTL heritability with RILs was approximately 9% for the three traits, assuming QTL heritability of 8% and 400 individuals assessed in generation 10r (12.5% greater than the heritability at generation 10r). Due to sampling, the estimated LD was greater with RILs than with non-inbred plants in generation 10r (Figure 6). Based on simulation 1, the average estimated Δ and r^2^ values were 0.0252 and 0.0241 for the RILs and 0.0235 and 0.0225 for generation 10r, respectively. Although these average values are equivalent, the estimated Δ values with RILs were four times greater on average than the estimated Δ values in generation 10r. Once again, the GWAS was ineffective when assuming low heritability and reduced sample size.

**Figure 6.**
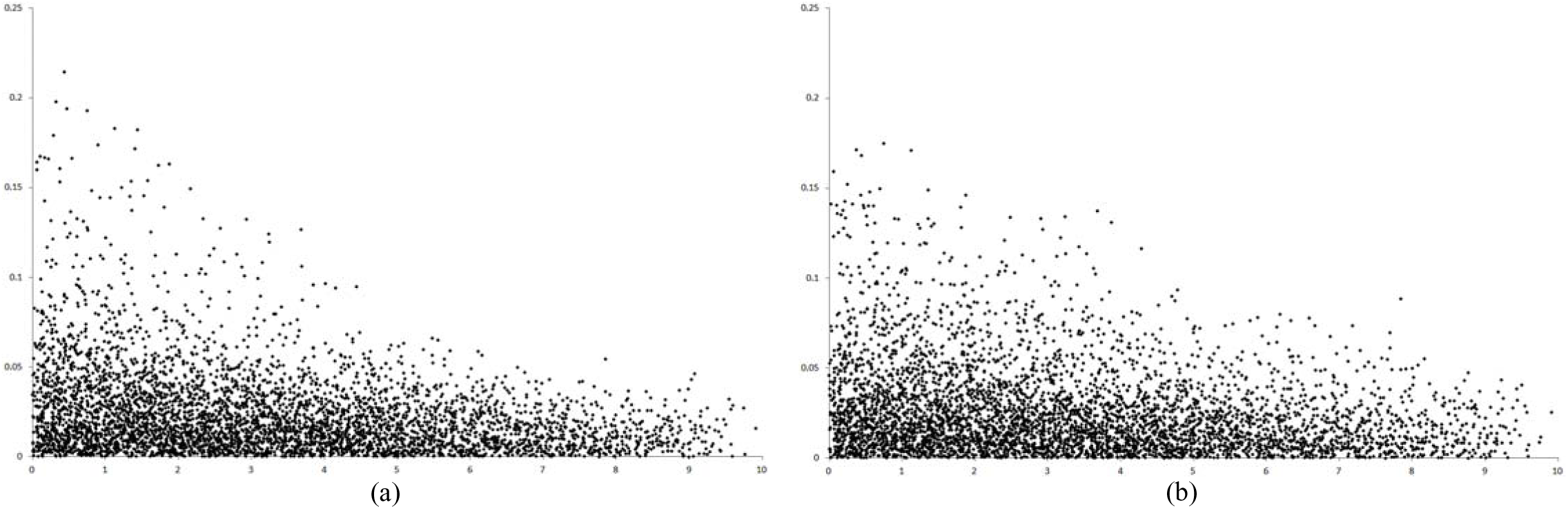
Relationship between the estimated LD value (absolute value; Y axe) and distance (cM; X axe) in population 1, generations 10r (random cross) (a) and 10r10s (random cross and selfing) (b), simulation 1, assuming a segment of 10 cM of chromosome 1 (centered on QTL 3).

## DISCUSSION

The presented theory proves that a significant association from a GWAS in a non-inbred or inbred open-pollinated population and in an inbred lines panel, while controlling the type I error rate and correcting for population structure and relatedness, is due to LD between the SNP and one or more linked QTLs. The theory also shows that GWAS provides estimation of the average effect of a SNP substitution (and consequently the estimation of SNP effects). Schaefer and Bernardo (2013) estimated SNP effects for days to anthesis, days to silking, oil and starch concentration, and measures of disease resistance using a maize inbred lines panel. We showed that only if there is a single QTL in LD with a significant SNP it is adequate to test dominance for the QTL loci. It is important to highlight that only if there is a single QTL in LD with a significant SNP, if the SNP is within the QTL, and if QTL and SNP alleles have the same frequency it is adequate to consider the SNP average effect of substitution as the QTL average effect of substitution. Furthermore, we also proved that a significant association due to admixture LD (population structure) does not depend on linkage disequilibrium between the SNP and a linked QTL.

To our knowledge, this is the first study on GWAS efficiency in open-pollinated population. The results are very encouraging and show that the process can be highly efficient, depending on LD, sample size, and QTL effect. In an open-pollinated population, the LD measure depends also on the SNP and QTL allele frequencies. Thus, significant associations involving several SNPs with the same QTL can be observed, including SNPs that are tens of mega base pairs (or centiMorgans) from the QTL. In reality, a closely linked QTL and SNP can have a lower LD value compared to a more distant QTL and SNP pair. In populations with low levels of LD, significant associations are expected to occur for only SNPs within the QTL or located very close to the QTL (within a few hundred base pairs), which favors the identification of a candidate gene for the QTL. In this scenario, a QTL would be declared based on one to a small number of significant associations spanning a chromosome region of a few kilo base pairs (not mega base pairs or centiMorgans).

A genome-wide association study is ineffective for lower sample size (200 individuals) and QTL heritability (4%), regardless of the population, i.e., including inbred lines panel and RILs. This scenario does not improve when increasing the FDR. Thus, we recommend that breeders employ larger sample size (400 individuals) and achieve high trait heritability (70-80%) (such as by genotyping parents and phenotyping replicated progeny). With intermediate (8%) to high (12%) QTL heritability and larger sample size, it is important to define a FDR of 1% to decrease the number of false-positive associations (note that some associations cannot really be false-positives). According to Larsson *et al.* (2013), false-positive associations can arise from markers that are in long-range LD with causative polymorphisms, which despite being rare are typically unaccounted for in association studies.

Our results are comparable to those obtained by Yu *et al.* (2008) in a simulation study investigating the genetic and statistical properties (power of QTL detection and FDR) of the nested association mapping (NAM) design. With 5,000 genotypes, they achieved an average power of QTL detection of 57% (with a range of 30 to 85%) when considering two trait heritabilities (0.4 and 0.7) and two different numbers of QTL controlling the trait (20 and 50). They also observed that a higher heritability always gave higher QTL detection power, particularly for QTL with moderate to small effect. Hung *et al.* (2012) assessed the maize NAM population and achieved heritabilities greater than 0.8 for traits related to flowering time and plant architecture, resulting in a good power to detect QTL. In contrast, traits with lower heritabilities (up to 0.6) and stronger sensitivity to environmental variation allow only a reasonable power of QTL detection. Also using the NAM population, Kump *et al.* (2011) evaluated resistance to southern leaf blight (SLB) disease and obtained a heritability of 87%. They identified 32 QTLs with predominantly small and additive effects on SLB resistance and many of the SNPs within and outside of QTL intervals were also within or near to genes previously shown to be involved in plant disease resistance.

Field results have demonstrated that GWAS are best carried out with a large sample size (Yu and Buckler 2006). According to Flint-Garcia *et al.* (2005), increasing the population size increases the number of individuals with rare alleles, thus improving the power to test the association between these rare alleles and the trait of interest. Yu *et al.* (2008) showed that the gain in efficiency by increasing sample size was evidenced by increased power of QTL detection and smaller FDR, mainly with heritability of 0.7 in comparison with a heritability of 0.4. Based on a simulation study, Long and Langley (1999) demonstrated that approximately 500 individuals should be genotyped for 20 SNP loci within the candidate gene region to detect marker-trait associations for QTLs that account for as little as 5% of the phenotypic variation. They observed that more power was achieved by increasing the population size than by increasing the SNP density within the candidate gene.

Compared to QTL mapping, GWAS is much more precise for mapping QTLs and identifying candidate genes. In QTL mapping studies based on simulated data, the bias in QTL position ranged from 2.0 to 6.0 cM depending on sample size, heritability, and marker density (Li *et al.* 2010). The bias with GWAS should be much lower because the significant SNPs are frequently within or very close to the candidate genes.

Compared to GWAS in an inbred lines panel, GWAS in open-pollinated population was less efficient, i.e., showed slightly lower power of QTL detection, higher number of false-positive associations, slightly higher bias in QTL position, and higher number of significant associations for the same QTL. The increase in efficiency by using the inbred lines panel was due to the lower degree of LD achieved by mixing groups of inbreds with positive and negative LD values. This is probably the main advantage of GWAS based on inbred lines panel. In contrast, when fixing the FDR, GWAS in a non-inbred population tends to be more efficient than GWAS with RILs from the same population. According to Flint-Garcia *et al.* (2005), the inbred lines panel exploits the rapid breakdown of LD in diverse maize lines, enabling very high resolution for QTL mapping. Population structure results from constructing a panel with inbreds from various breeding programs and distinct heterotic groups, which can cause false-positive marker-trait associations if the data is not corrected (Yan *et al.* 2009). The lowest parametric LD values for the inbred lines panel occurred in published studies (Yan *et al.* 2009, Remington *et al.* 2001). Moreover, with the inbred lines panel, generally, only SNP loci within the QTL showed significant association, which is a highlighted result from GWAS that can serve as a basis for a fine mapping strategy for marker-assisted selection and map-based cloning genes (Gupta *et al.* 2005).

GWAS in plant breeding has been effective for identifying candidate genes for quantitative traits such as plant architecture, kernel composition, root development, flowering time, drought tolerance, pathogen resistance, and metabolic processes (Zhu *et al.* 2008). Based on our evidence, breeders can employ non-inbred and inbred breeding populations while taking into account that the level of LD should be low, the sample size should be higher than that necessary for QTL mapping, and the QTL heritability should be intermediate to high to achieve greater power of QTL detection and precise mapping of candidate genes.

## ACKNOWLEDGMENTS

We thank the National Council for Scientific and Technological Development (CNPq), the Brazilian Federal Agency for Support and Evaluation of Graduate Education (Capes), and the Foundation for Research Support of Minas Gerais State (Fapemig) for financial support.

